# Signatures of selection and mechanisms of insecticide resistance in Ugandan *Anopheles funestus*: Insights from embedding translational genomics into the LLINEUP cluster randomised trial

**DOI:** 10.1101/2025.06.17.659961

**Authors:** Lilian Namuli-Kayondo, Sanjay C Nagi, Harun Njoroge Nganga, Anastasia Hernandez-Koutoucheva, Daniel P. McDermott, Samuel Gonahasa, Amy Lynd, Ambrose Oruni, Catherine Maiteki-Sebuguzi, Jimmy Opigo, Adoke Yeka, Agaba Katureebe, Mary Kyohere, Moses R Kamya, Grant Dorsey, Janet Hemingway, Sarah G Staedke, Samuel L Nsobya, Joaniter I Nankabirwa, Jonathan Kayondo, Chris Clarkson, Alistair Miles, Mara K N Lawniczak, Eric R Lucas, Martin J Donnelly

## Abstract

In response to the emerging threat of insecticide resistance in malaria vectors, insecticides are being repurposed for vector control or developed *de novo*. Good stewardship of these finite new resources is essential if disease control programmes are to remain effective. This is dependent on timely data to help guide evidence-based decision-making for National Malaria Control programmes (NMCPs). By embedding genomics into cluster randomized control trials (cRCTs), we can perform surveillance and early detection of insecticide resistance variants to new and repurposed chemicals in natural field conditions, supporting effective stewardship.

The LLIN Evaluation Uganda Project (LLINEUP) trial evaluated the efficacy of pyrethroid-piperonyl butoxide (PBO) and pyrethroids-only long-lasting insecticidal nets (LLINs). It was conducted in Uganda between 2017-2020 and was the largest cRCT to date, covering 40% of the country in 104 health sub-districts. We embedded genomic surveillance within LLINEUP to detect and track insecticide resistance variants. At baseline and throughout the trial, we sampled *Anopheles* mosquitoes with Prokopack aspirators and performed Illumina whole-genome sequencing.

We show that *An. funestus* populations were relatively unaffected by the interventions, compared to *An. gambiae* s.l., which were markedly reduced six months following LLIN deployment. Standard approaches for describing genetic diversity and population structure e.g. fixation index (*F*_*ST*_), Principal Component Analysis (PCA) and Neighbour-Joining (NJ) trees, were consistent with the density observations and suggestive of a single large *An. funestus* population in Uganda with little genetic differentiation. Genome-wide selection scans revealed strong signals of selection at the *Resistance to pyrethroid-1 (RP1*) locus and *Cyp9k1*, both loci previously implicated in pyrethroid resistance. We report two additional loci, eye diacylglycerol kinase (*Dgk*) (≅13.5Mb on the X chromosome) and O-mannosyl-transferase (*TMC-like*) (≅67.9Mb on 3RL) that showed signals of selection. Known DDT and permethrin resistance-associated variants at the *Gste2* locus, L119F and L119V, were also identified. Over the trial period, changes in haplotype frequencies were observed in regions under selection, with more pronounced shifts in the PBO arm. Notably, there were significant reductions in the frequencies of swept haplotypes (measured by delta (Δ) H12) in the *Dgk* and *Cyp6p9a* regions, while significant increases in haplotype frequency were observed at *Gste2* and *Cyp9k1* loci.

Our findings reveal the differential impact of the trial on *An. gambiae* s.l. and *An. funestus* densities and the differing responses of *An. funestus* populations to pyrethroid and pyrethroid-PBO selection pressure. These insights underscore the potential value of tailored, species- and region-specific vector control strategies, supported by regional genetic surveillance, to better control insecticide resistance evolution and spread. By embedding genomic surveillance in cRCTs we can facilitate the discovery of putative resistance variants and can provide evidence of their impact on vector control tool efficacy; both of crucial importance to evidence-based deployment of vector control tools by NMCPs.

## Introduction

Malaria vector control and elimination efforts remain dependent on the use of LLINs and indoor residual spraying (IRS). In sub-Saharan Africa, LLINs averted an estimated 68% of malaria infections between 2000 and 2015^1^ . Vector control, however, is facing notable challenges, such as shifts in behaviour and the species composition of malaria vectors^2^, and the emergence and spread of insecticide resistance^3,4^. With insecticide resistance now widespread in many African malaria-endemic countries, the future of chemical vector control efforts is of critical concern^5^. Uganda, a country that accounted for the third highest number of malaria cases in Africa in 2023^3^, is experiencing changes in Anopheles mosquito populations and widespread insecticide resistance following deployment of LLINs and IRS^2,4,6,7^. We have previously documented increases in the frequency of genetic markers associated with pyrethroid resistance in *An. gambiae* s.l. across Uganda^4^ whilst in Tororo district in eastern Uganda, an area historically dominated by *An. gambiae* s.s., shifts to *An. arabiensis* and subsequently to *An. funestus* s.s., as the predominant vectors, have been observed following rounds of IRS with different insecticides^7^. Studies in Uganda suggest resistance to pyrethroids and dichloro-diphenyl-trichloroethane (DDT) is widespread^6,8,9^. A robust and real-time resistance surveillance system is needed if we are to detect and track resistance, especially to new insecticides, in real-world settings.

The LLINEUP trial was a cluster-randomised trial designed to compare the efficacy of new generation pyrethroids-PBO LLINs to pyrethroid-only LLINs. The trial was conducted between 2017 and 2020 in 104 health sub-districts (clusters) in 48 districts, covering 40% of the country^10^. The study demonstrated that pyrethroid-PBO LLINs were associated with a significant reduction in mosquito densities and malaria prevalence among children aged 2-10 years, compared to pyrethroid-only LLINs, over 2 years of follow-up^11^. Overall, in the LLINEUP trial, *An. funestus* was found in 63 of the 104 clusters and was predominant in 23 clusters^4^. LLINEUP offered us an opportunity to study the impact of interventions on vector densities, population structure, and to track the emergence and spread of insecticide-resistance mutations in natural conditions.

Here, we use whole genome sequencing (WGS) data from mosquitoes collected in the LLINEUP trial to understand how vector populations respond to insecticide pressure when subject to mass distribution of LLINs aimed at achieving universal coverage. Recent studies have shown apparent changes in *An. funestus* behaviour with an increased propensity to rest and bite outdoors and during the daytime^12^, and higher *Plasmodium* infection prevalences than *An. gambiae* s.s.^13^ Moreover, *An. funestus* populations are becoming increasingly resistant to pyrethroids^8,14^, with experimental hut work showing that resistant strains may reduce the efficacy of treated nets^15^. In this paper we focus upon trial impacts upon *An. funestus*, while a companion paper Njoroge *et al*^*16*^ focuses on *An. gambiae* s.s. Utilizing WGS data, we tested the hypothesis that the use of pyrethroid-PBO LLINs would exert a directional selection pressure on *An. funestus* s.s *(henceforth An. funestus)*; and that the apparent rebound in vector abundance seen in the trial is mirrored to changes in population diversity.

## Results

### Hypothesis 1: The use of LLINs results in a decrease in both Anopheles spp. vector density

Maiteki et al showed that, during the LLINEUP trial, there was a sharp decline in *Anopheles spp* vector densities following LLIN deployment^11^. We tested the hypothesis that there was a reduction in both *An. gambiae* s.l. and *An. funestus* s,l. densities following LLIN deployment. We disaggregated the data for the two major vectors and showed that whilst LLIN use was followed by a significant reduction of *An. gambiae* s.l. vector densities (p=0.0216), *An. funestus* was seemingly less affected by the interventions (p=0.253) (Fig. 1 B&C and supplementary Table 1&2).

**Fig 1:**
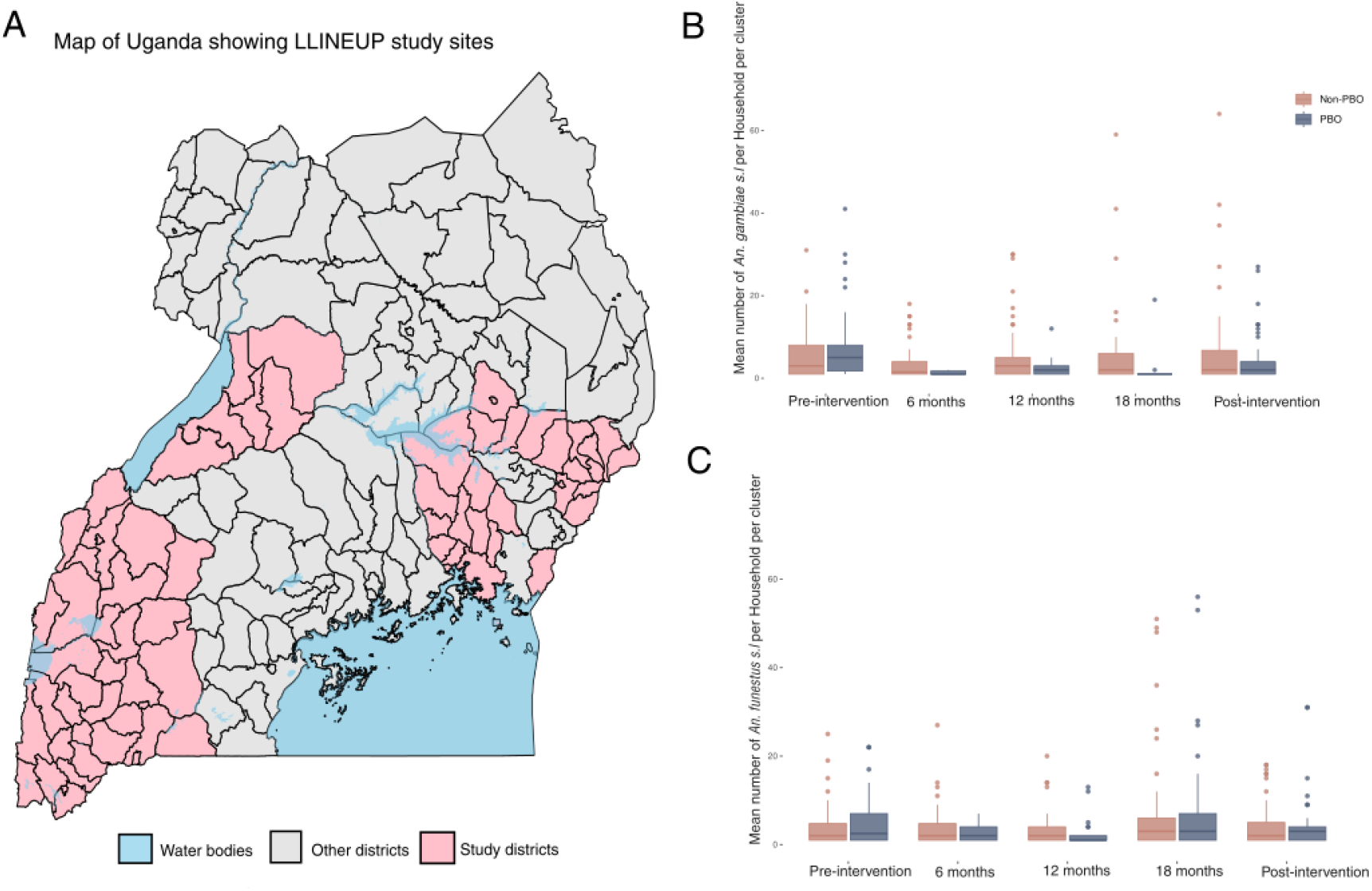
A: Map of Uganda showing LLINEUP sample collection sites. (pink) and the rest of the districts in the country (grey area). Box-whisker plots of numbers of <u>An. gambiae s.l.</u> (B) and <u>An. funestus</u> (C) per household during the trial. The Y-axis is truncated to 75 to improve visual clarity. We performed a Negative Binomial Model (glmmTMB (species∼ round+LLINtype+ (1|household), family=nbion2) to determine the number of vectors per household.

### Hypothesis 2: The Eastern and Western Ugandan An. funestus populations are genetically structured

The East and West Uganda LLINEUP sites are geographically discrete. Given there was a marked discontinuity between collections from the two regions, (Fig. 1), we tested for evidence of population structure using the 60-80Mb region of chromosome 2, in which known chromosomal inversions are absent. We first examined evidence of population structure in the context of other *An. funestus* populations from the *An. funestus* 1000 genomes project https://www.malariagen.net/project/anopheles-funestus-genomic-surveillance-project/. Our analysis showed that, consistent with the population structure described by Bodde et al (2024)^17^, the Ugandan LLINEUP population grouped with the equatorial cluster (Supplementary Fig. 1). When analyzed independently of the Af1000 samples, the Ugandan *An. funestus* populations, despite being collected from non-contiguous sites, did not show evidence of marked spatial or temporal population structure, suggesting a single, genetically connected population (Fig. 2A and Supplementary Fig. 2). We further investigated possible genetic differentiation due to their geographical separation using *F*^*ST*^ statistics. Both the within and between region pairwise *F*^*ST*^ values were consistently very low (≤0.003), suggesting that the Ugandan *An. funestus* is one genetically connected population (supplementary Table 3). We therefore considered the Ugandan *An. funestus* as a single cohort for all further analysis.

**Fig 2:**
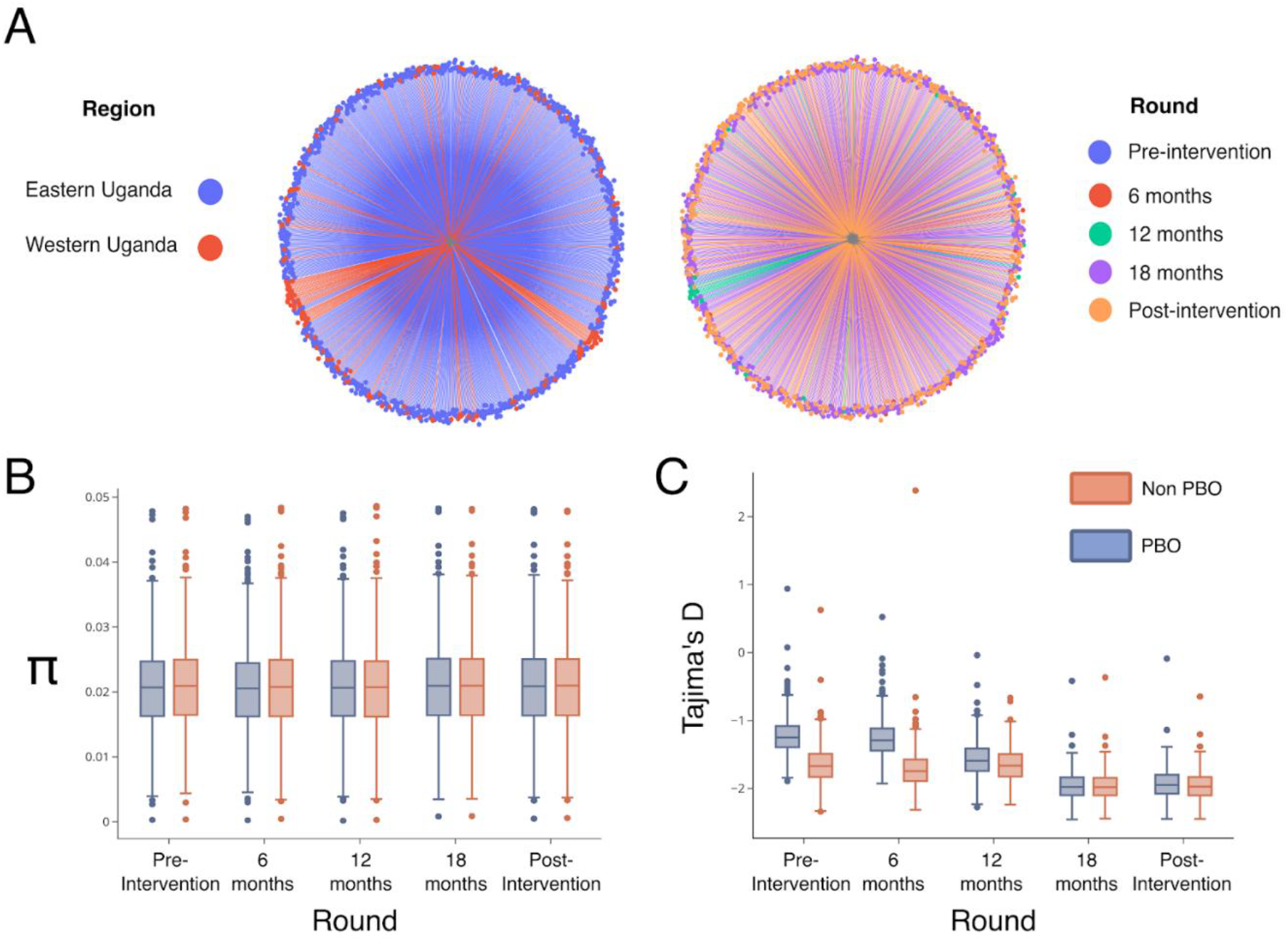
Population structure and genetic diversity. A) Neighbour-joining trees (NJT) showing a lack of population structure in <u>An. funestus</u> population from Eastern and Western Uganda (left) and during the different trial rounds (right). NJTs, B) Nucleotide diversity (π) and C) Tajima’s D were calculated using the genomic region 2R:60-80Mbps to avoid polymorphic chromosomal inversions.

### Hypothesis 3: *The use of PBO LLINs leads to a population bottleneck*

Based on a demographic expectation that LLIN deployment would lower mosquito densities, we hypothesized a resultant population bottleneck that would reduce genetic diversity through inbreeding-driven loss of rare alleles. We determined changes in *An. funestus* genetic diversity over the course of the intervention in each of the intervention arms. Both nucleotide diversity (π) and Tajima’s D maintained a broadly uniform pattern during the trial. Tajima’s D was consistently negative, with post-intervention samples more negative than baseline (pre-intervention) samples, supporting a possible population expansion (Fig. 2B and C).

### Hypothesis 4: The use of LLINs exerts selection pressure on mosquito vectors that increases the prevalence of insecticide resistance-associated alleles

We performed a genome-wide single nucleotide polymorphism (SNP) association study to look for trial-specific changes in SNP frequencies over time. After corrections for multiple testing, no significant differences in SNP frequencies were observed between pre- and post-intervention mosquito populations (Supplementary Fig. 3).

We analysed haplotypes using Garud’s H12 (Fig. 3A) to identify regions under recent positive selection, and delta (Δ) H12 (Fig. 3B) to identify H12 signals that changed in frequency during the trial. For loci that were detected to be under positive selection, a model-based approach was employed to pinpoint corresponding genomic coordinates. Using signal intensity and spatial closeness, high-scoring loci were grouped into distinct peaks, and regions above a predetermined H12 threshold (top 1% genome-wide) were identified (Supplementary Table 4). Five regions showed signatures of recent selection; of these, two regions containing known resistance-associated loci *Cyp9k1* (X chromosome-8Mb) and *RP1 (resistance to pyrethroids) locus* (2 chromosome-8Mb) had stronger H12 signals. Three other regions; the 2RL chromosome at 76Mb, 3RL chromosome at 67Mb, and the X chromosome at 13.5Mb, showed weaker signals of selection (Fig. 3A). These signals correspond to Glutathione S-transferase Epsilon (*Gste2*) and Protein-O-mannosyltransferase Transmembrane channel-like protein and Eye diacylglycerol kinase (*Dgk*). While the *Gste2* gene has been implicated in resistance to both DDT and permethrin^18^, Protein-O-mannosyltransferase Transmembrane channel-like (*TMC-like*) protein (*henceforth* TMC) is a ‘novel’ gene in *Anopheles* insecticide resistance profile. In *Caenorhabditis elegans*, the *TMC* gene has been linked to alkali and sodium sensing, enabling detection and avoidance of noxious alkaline environments ^19,20^. Worms with a mutated version of the gene maintain egg-laying even in harsh conditions ^21^. In contrast, the *Dgk* gene has recently been found to be under positive selection in *An. gambiae* s.s.^*16,22*^, however, its role in resistance is yet to be determined. In *Drosophila* ^*23*^, the gene is reported to play a role in light sensitivity and has also been reported to function in the regulation of acetylcholinesterase in nerve impulse transmission in worms ^24^. For both the ‘novel’ *TMC* and *Dgk* loci, we further investigated whether the haplotypes driving the selection signals were shared between East African populations across space and time. We found that haplotypes potentially driving selective sweeps at these loci were present in the Uganda mosquito populations as early as 2014 (Supplementary Fig. 4). Haplotypes at both *Dgk* and *TMC* loci appeared to be shared between Uganda, Kenya, and Tanzania populations (Fig. 4A&B). We did not see any evidence of selection in the *Vgsc* gene, as previously reported in a study of *An. funestus* populations in neighbouring Tanzania^25^.

**Fig 3.**
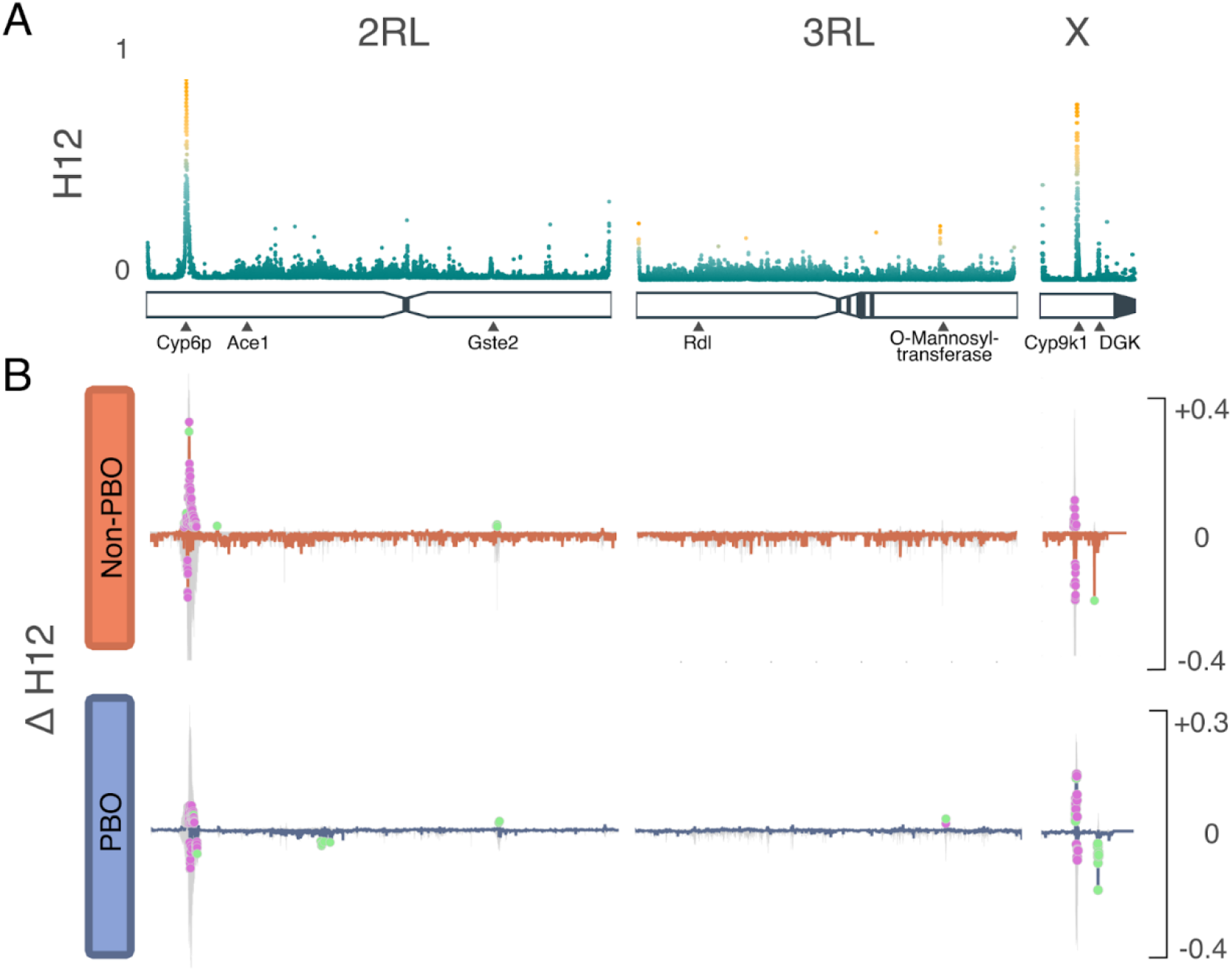
Selection analyses. A) H12 genome-wide selection scans of the 2RL, 3RL, and X chromosomes. Scans were performed using 500 SNP windows; peaks are labeled by loci likely to be driving resistance. Increase haplotype frequency; on the 2RL chromosome, the signal is centered ∼8,600,000Mb corresponding to RP1 locus and another signal ∼76,400,000 corresponding to the Glutathione S-transferase epsilon 2 gene; on 3RL chromosome, the signal centered ∼67,900,000 corresponding to O-Mannosyl-transferase; on the X chromosome, signal at ∼8,400,000Mb corresponding to Cyp9k1 gene and another signal ∼13,004,000 corresponding to the Diacylglycerol Kinase gene. B) ΔH12 analysis of the 2RL, 3RL, and X chromosomes. Change in haplotype frequency (ΔH12) in the Cyp6p9a, and Cyp9k1 (2 and X chromosomes), a decrease in haplotype frequencies in the Dgk region on the X chromosome. The colored dots show windows that surpassed permutation-based significance threshold (purple=marginal; green=significant).

**Fig 4:**
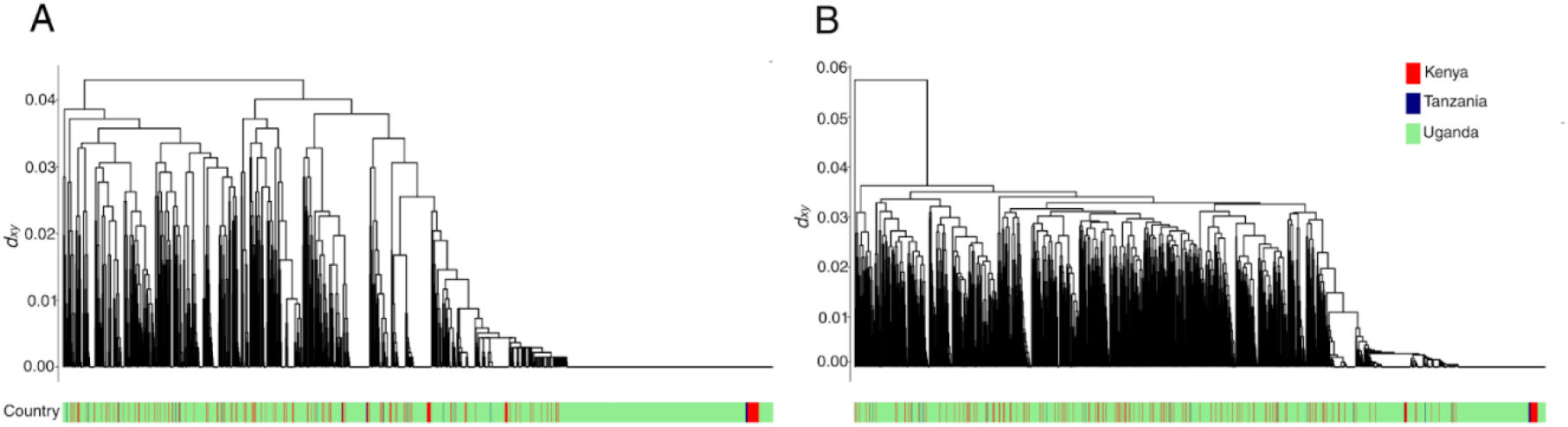
Hierarchical clustering. dendrogram of haplotypes over the A) TMC (3RL: 67,925,771-67,931,656 and B) Dgk (X: 13,590,000-13,690,044) genes. Dendrogram leaves are labeled by country of mosquito population origin.

To determine temporal changes in the swept haplotypes during the trial period, we calculated ΔH12, the difference in H12 values between pre- and post-intervention populations. Negative ΔH12 values are indicative of increasing haplotype diversity, with positive ΔH12 values indicating reduced haplotype diversity, suggestive of increased selective pressure following the intervention. For each chromosome arm, pre-intervention and post-intervention H12 were calculated in 500bp genomic windows. For swept haplotypes, we defined peaks as windows with ΔH12 more than three times the difference between the median and the 98th percentile of the genome-wide ΔH12 distribution. To determine the statistical significance of a change in ΔH12, we generated 1000 random permutations of the labels for the pre- and post-intervention phases. We then recalculated ΔH12 for each window under the null distribution. We obtained p-values by comparing permutated values to the observed ΔH12 using a two-sided test. Genomic windows

were considered significant if p<0.01. The ΔH12 (Fig. 3B) was plotted as the difference in the pre- and post-intervention, highlighting green regions with significant change in H12 signal. Whilst we observed an increase in the frequency of swept haplotypes in the *RP1* locus in the Non-PBO arm, haplotype frequency at the loci decreased in the PBO arm, which also exhibited an increase in haplotypes at *Cyp9k1* and *TMC* loci. The *Dgk* signal decreased in frequency in both study arms, but the reduction was arguably more pronounced in the PBO arm. There was reduced haplotype diversity at the *Gste2* loci in both trial arms.

To further examine genomic regions under positive selection, we carried out diplotype clustering to identify single-nucleotide polymorphism (SNP) mutations that may be driving selective sweeps^26^. Both the novel *TMC (*Supplementary Fig. *4A)* and *Dgk* (Supplementary Fig. 4B) loci showed signals consistent with a major sweep with haplotypes that appear to be shared between Eastern and Western Uganda populations. At the *Dgk* locus, individuals with reduced heterozygosity appeared to mainly carry the A339V SNP mutation. Like previous work, our findings show that, in the Ugandan cohorts, the *Cyp9k1*-G454A mutation was associated with a hard selective sweep^17,27^ (Supplementary Fig. 5D). Mutations in the *Cyp9k1* and the *Cyp6p9a* mutations have been implicated in pyrethroid resistance^28^ and are suggested to have pleiotropic effects beyond resistance, potentially influencing fitness-related benefits^17^. In the *Gste2* gene cluster, besides the previously reported L119F mutation associated with pyrethroid resistance, we observe three further SNPs, L119V, L175P and D40E (Supplementary Fig. 5B).

## Discussion

### Population structure and diversity of Ugandan An. funestus

Understanding vector population demographic history provides valuable insights into the likely response of vector populations to insecticide pressure^17^ and the likelihood of the spread of adaptively advantageous variants^29^. We demonstrate that while the trial reported a significant decline in *Anopheles* vector densities^11,30^, this was mainly due to changes in *An. gambiae* collection numbers and that *An. funestus* abundance was less affected by LLIN use. A corollary of this observation comes from the nucleotide diversity data (Fig. 2), which were stable throughout the trial. However, it should be noted that diversity indices are slow to change in organisms with large effective population sizes^31^. Our results are contrary to previous studies that have reported population bottlenecks and reduced genetic diversity in other *Anopheles* vectors following vector control^32-34^.

We observed that *An. funestus* in Uganda is best described as a panmictic population with limited evidence for genetic differentiation. This suggests that despite substantial spatial separation (several hundred kilometers), the Ugandan *An. funestus* may be considered a genetically homogeneous, unstructured population with high gene flow. Previous studies in Uganda using microsatellite markers have reported the role of geographical distance on the population structure and genetic differentiation of *An. funestus s*.*s*^*35-37*^. In contrast, our study focused on a 20Mb region free of known inversions, using individual whole genomes to examine genetic structure. While we did not explicitly asses the effect of geographical distance on population structure, differences in reported genetic structure may be due to marker type and genomic regions analysed. Inversions and microsatellites have been reported to show signals of natural selection in *An. funestus*, as they are often influenced by linkage to nearby adaptive loci^36,38^. Thus, the two markers may not provide a reliable neutral population structure and differentiation compared to a naturally recombining region. Our findings are indicative of genetically homogeneous *Anopheles funestus* populations with possible high gene flow. In regions where *Anopheles* populations are genetically connected, insecticide resistance mutations rising due to selection in one region may disperse widely via gene flow^35,39^. Dispersal of adaptive alleles possess a major obstacle to ongoing vector control programs, potentially undermining the future efficacy of interventions. To mitigate unintended cross-regional insecticide resistance spread, a proactive country wide resistance management strategy coordinated through the National Malaria Elimination Program is essential to preserve the efficacy of the current and future vector control tool kit.

### Evidence of metabolic resistance

We had an opportunity to monitor the evolution and spread of insecticide resistance variants in *An. funestus*, and how these are sustained in natural field conditions during LLIN use. In *An. gambiae* s.l, and to a lesser extent in *An. funestus*, WGS data has been used to elucidate both known and ‘novel’ mutations involved in resistance^40^. Additionally, the use of H12 statistics has revealed regions of the vector genome that are under selection, unveiling mechanisms that drive resistance in wild populations. Using the same approach, we examined *An. funestus* genomes to identify regions under positive selection and pinpoint both known and novel loci under selection. Insecticide resistance in *An. funestus* has been reported to be mainly driven by metabolic enzymes, primarily cytochrome P450s. We identified strong recent selection signals in two gene clusters *Cyp6p9a* and *Cyp9k1*, previously associated with metabolic resistance in Uganda *An. funestus*^*27,35,41*^. Furthermore, we show that the sweep in Cyp9k1 is tagged by the SNP variant *Cyp9k1-*G454A as previously reported^27^. These findings reinforce the existing body of evidence that resistance within this species is mainly driven by metabolic genes. Evidence of the importance of cytochromes and *Gste2* in *An. funestus* metabolic resistance to pyrethroids has been previously reported^29^; phenotypically resistant *An. funestus* had higher expression of the genes compared to susceptible strains^28^. Comparably, in *An. gambiae* s.l SNP mutations and copy number variants in cytochromes and *Gste2* have been strongly associated with resistance to pyrethroids and DDT^42,43^.

### LLIN-specific changes

The ΔH12 statistic was used as a proxy for studying LLIN-specific changes in haplotype frequencies of genomic regions under selection during the trial^44^. We observed significant changes in haplotype frequencies in five major genomic regions; *RP1 locus, Gste2, TMC, Cyp9k1*, and *Dgk*. More significant changes in haplotype frequencies were observed in the PBO arm. Particularly, there was an increase in *TMC* and *Cyp9k1* haplotype frequencies. Both TMC and *Dgk* loci are novel in *An. funestus* insecticide resistance profile. In worms, the *TMC* locus has been shown to play a role in the evasion of alkaline and noxious environments^19^ and could play a similar role in insects. A significant reduction in *Dgk-swept* haplotype frequencies was observed in both trial arms, albeit more pronounced in the PBO clusters. Although the gene has been previously reported as under selection in *An. gambiae*, its role is yet to be elucidated. We report that the major sweeps in both novel loci were shared among populations in Uganda, Kenya, and Tanzania, and that these sweeps were present in the Uganda populations sampled in 2014. The presence of shared haplotypes in East African populations points to either common ancestry or gene flow, while the presence of the sweep in 2014 Ugandan samples highlights the potential relevance of the two loci in local vector populations. Selection pressure around the *Gste2* gene cluster was observed in both trial arms. Consistent with another LLINEUP study using molecular markers^4^, we report counterintuitive shifts in haplotype frequencies in characterised cytochrome loci. Although we observed a decrease in haplotype frequencies in the *RP1* locus in the PBO trial arm, the detected increase in *Cyp9k1* haplotype frequencies in the arm was rather unexpected given the anticipated inhibitory effects of PBO on cytochrome expression. We postulate that in presence of PBO, selection shifts from the *RP1* loci and instead acts on the alternative loci *Cyp9k1*. It is plausible that *Cyp9k1* represents an alternative detoxification pathway less susceptible to PBO inhibition and is more strongly selected for when mosquitoes are exposed to PBO based interventions. Moreover, the *TMC* locus may reflect novel or compensatory resistance mechanism that is conceivably activated under PBO induced insecticide stress. The *Gste2* loci, on the other hand, appears to function as a broad-spectrum detoxification pathway and potentially plays a role in both pyrethroid and PBO-associated detoxification. *Anopheles funestus* with *Gste*-L119F mutation has been implied to have an increased survivorship when exposed to PBO-pyrethroid nets^15,45^. Moreover, mosquitoes with increased pyrethroid intensity resistance did not exhibit significant differences in their expression levels of *Cyp6p9a* and *Cyp9k1* genes^28^. The differential selection of the metabolic gene loci in the two trial arms needs further investigation.

## Conclusion

This study demonstrates the value of embedding genomic surveillance within cluster randomised trials. We show that *An. funestus* populations in Uganda remained resilient to interventions during the LLINEUP trial, with multiple loci in the genome under selection. The observed increase in diversity at *RP1* locus and concurrent decrease in diversity at the *Cyp9k1* and the novel *TMC* loci under PBO pressure suggests a potential shift of resistance pathways in *An. funestus* when classical enzymes are inhibited. Overall, the genetic connectivity observed in the vector highlights possible transregional spread of resistance alleles. These findings underscore the role of tailored vector control strategies, continuous genomic surveillance, and regionally informed insecticide deployment. Embedding genomic surveillance within cluster randomized trials enables the detection of emerging resistance mechanisms and provides real time data to inform evidence-based vector control strategies.

## Methods

### Sample collections

The Long-Lasting Insecticidal Net Evaluation in Uganda Project (LLINEUP) (Staedke 2019), was a cluster randomised control trial in Uganda in 2017-2019, that compared pyrethroid LLINs with and without the synergist PBO. The LLINEUP was the largest bed-net trial to date, covering 40% of the country and involving 104 clusters (health sub-districts), and 48 districts (38 clusters in eastern and 66 clusters in western Uganda). Clusters were randomised to receive either standard or PBO LLINs^10^. As outlined by Amy Lynd et al. in 2019, one household (of the 50 included in the community cross-sectional survey) with a child between 2-10 years was randomly selected for entomological survey collections. Morning Prokopack aspiration collections, conducted by a single individual for 10 minutes before 10:00 hrs in each household, were used to collect mosquitoes^46^. Mosquitoes were counted and morphologically identified up to the genus level. For each mosquito, DNA was extracted using a Nexttec DNA extraction kit (Biotechnologie GmbH), and samples were amplified for *An. funestus* sl sibling species identification, using the Koekemoer et al., 2002 protocol^47^.

### Changes in vector abundance during the study

Separate analysis for *An. funestus* and *An. gambiae* s.l. were conducted to assess the impact of net type on temporal trends of mosquito densities. For each species we filtered data according to sibling species, and then grouped it by household, enumeration area, survey round and net type. Exploratory analysis using boxplots and histograms was done to visualize density distributions by net type during the different survey rounds. Because the data was over dispersed, we used a generalized linear mixed model (GLMM) with a negative binomial distribution, with net type and survey round as fixed effects and households as random effects. The model was fitted using the glmmTMB package.

### Whole genome sequencing and bioinformatics analysis

Using the Illumina HiSeq X platform, we performed sequencing at the Wellcome Sanger Institute as part of the MalariaGen/Vector Observatory projects following the pipeline (pipeline full details here: *https://malariagen.github.io/vector-data/ag3/methods.html*). we then analysed a total of 1150 whole genome sequences and of these, 954 were from Eastern and 196 from Western Uganda (Supplementary Table 4)

Briefly, following the manufacturer’s instructions, paired-end multiplex libraries (each multiplex containing 12 individually tagged mosquitoes) were created. Cluster generation and sequencing were carried out according to the manufacturer’s instructions. To minimise fluctuation in yield, between sequencing runs three sequencing lanes were generated. Sequencing was performed using paired-end 100-bp sequence reads, with the insert size ranging from 100 to 200 bp. Sequence reads were then aligned to the AFunGA1 *An. funestus s*.*s*. reference genome, using the BWA algorithm. Following recommended best practices, SNPs were identified using the GATK tool version 3.7-0 RealignerTargetCreator and IndelRealigner. Additionally, where short reads could be mapped uniquely, and there was a minimal indication of structural variation, the generated alignments were used to identify genomic areas that were accessible for SNP calling. Statistical phasing of sequencing data was conducted using SHAPEIT2.

The *An. funestus* reference genome^48^ as well as genetic variants (haplotypes and SNPs), were accessed via the MalariaGEN repository while bioinformatics analysis was performed using both the R programming language^49^ and the malariagen_data python package^50^.

### Population structure

We assessed geographical population structure using NJTs supplemented with PCA. To determine if there was temporal and spatial gene flow between the Eastern and Western Uganda populations during the trial period, we computed pairwise Hudson’s F_*ST*_ and windowed genome-wide F_ST_ scans in the Eastern and Western Uganda cohorts across the different study rounds. F_ST_ is aimed at calculating genetic differentiation between two cohorts (population). Since selective sweeps and inversions in the genome could result in variations in alleles within the population, we used the 60Mb-80Mb on the 2RL (an area with no known inversion or selective sweeps), to determine levels of population structure and genetic differentiation measurements.

To estimate genetic differentiation between and within the East and West Ugandan populations, we calculated pairwise F_ST_ using Patterson’s estimator across 1,000 SNPs implemented in the *scikit-allel* python package. For within populations diversity, pairwise F_ST_ was determined for two randomly selected individuals from the same population while for between populations, diversity was determined by picking one randomly chosen individual from each of the populations and their pairwise F_ST_ computed. This process was repeated for 1,000 iterations to obtain a null distribution of F_ST_ values. Mann-Whitney U tests comparisons were used to compare distributions of the pairwise F_ST_. To determine genetic differentiation, within the two population F_ST_ values for East and Western populations were compared to the between-population F_ST_ values.

### Genetic diversity statistics

Population diversity was analysed using nucleotide diversity (theta-π), and Tajima’s D. Theta-pi measures the average number of pairwise differences between DNA sequences in a cohort; a higher value implies more nucleotide differences between the DNA sequences. Watterson estimator (theta-w); scaled by a constant, quantifies the number of segregating sites within a cohort with a high value, implying many segregating sites within the analysed genome. Tajima’s D is the difference between Theta-w and theta-π scaled to a constant; a negative value indicates the presence of rare alleles. A constantly growing population will have a negative Tajima’s D while a positive value represents the absence of rare alleles, possibly due to a recent population bottleneck <u>^39^.</u>

### Studying selective sweeps

To establish evidence of recent selection, we used Garud’s H12 statistic^51^, a statistic with the power to detect both soft and hard selective sweeps within populations. Genome-wide H12 scans were calculated in 500bp SNP windows. ΔH12 was calculated by subtracting the H12 measurements of round 5 from baseline H12 measurements^43^. For each peak (selective sweep) on the H12 selection scan, diplotype clustering analysis was done to determine SNPs, carried by the swept haplotypes; dendrogram leaves were shaded to represent individual samples. H1X genome-wide selection scans were done, to determine if the observed sweeps were shared over space and time.

### Diplotype clustering

Diplotypes from the beginning to the end of each of the four gene loci that were under selection: *RP1* (2RL: 8,685,464-8,692,407), *Cyp9k1* (X: 8,440,000-8,451,000), *Gste2* (2RL: 76,406,705-76,407,655), *TMC (3RL: 67,925,771-67,931,656)* and *Dgk* (X: 13,590,000-13,690,044) were obtained. Diplotypes were grouped using complete-linkage hierarchical clustering. The pairwise difference between this array of allele counts for each individual in the pair was determined for each site. The distance metric and full linkage were then determined by using the city-block (Manhattan) distance.

### Genome-wide association analysis

Following Lucus et al 2024, we performed SNP-wise GWAS using SNPs with no missing data<u>^42^.</u> For each SNP, to determine if there were any changes in SNP frequencies within the mosquito genome between baseline and round 5 of the trial, we fitted a binomial GLMM models (using the glmmTMB package) with trial rounds as a fixed effect and households as a random effect. We obtained a P-value of association for round for each SNP. False discovery rate test^52^ with adjusted p-values <0.05 was performed to control for multiple testing.

### Data availability

The LLINEUP trial was approved by the Liverpool School of Tropical Medicine (ref 16-072) and the Uganda National Council for Science and Technology (UNCST ref HS 2176. We present the analysis of the *An. funestus* sample genomic data is from the “1288-VO-UG-DONNELLY-VMF00219”. The LLINEUP genomic data is publicly available through the malariagen_data python API (https://malariagen.github.io/malariagen-data-python/latest/index.html). The scripts that were used for data processing, population genetics and figure generation in the current study are available publicly at the LSTM GitHub repository at https://github.com/vigg-lstm/LLINEUP-genomics.

### Funder information declared

National Institute of Allergy and Infectious Diseases of the National Institutes of Health, RO1AI116811; Bill & Melinda Gates Foundation, OPP1210750; Fogarty International Center of the National Institutes of Health, D4TW010526; Bill & Melinda gates Foundation, INV-069333.

