## Supplementary_Tables_Figures for "Signatures of selection and mechanisms of insecticide resistance in Ugandan *Anopheles funestus*: Insights from embedding translational genomics into the LLINEUP cluster randomised trial"

**Supplementary material**

Supplementary Table 1

| **Intervention arm** | **Pre-intervention** | | **6 months** | | **12 months** | | **18 months** | | **Post-intervention** | |
|---|---|---|---|---|---|---|---|---|---|---|
| | AF | AG | AF | AG | AF | AG | AF | AG | AF | AG |
| **Non-PBO** | 176 | 549 | 149 | 190 | 177 | 384 | 394 | 295 | 242 | 609 |
| **PBO** | 257 | 816 | 45 | 37 | 73 | 119 | 325 | 78 | 193 | 280 |

Table 1: Total number of PCR confirmed *An. gambiae* s.l. and *An. funestus* s.l. vector densities during the LLINEUP trial. AG= *Anopheles gambiae* s.l.; AF= *Anopheles funestus* s.l.

Supplementary Table 2

<table>
<tr><td></td><td colspan="2">Estimated</td><td colspan="2">Std error</td><td colspan="2">Z value</td><td colspan="2">Pr(>|z|)</td></tr>
<tr><td></td><td>AG</td><td>AF</td><td>AG</td><td>AF</td><td>AG</td><td>AF</td><td>AG</td><td>AF</td></tr>
<tr><td>Intercept</td><td>1.860</td><td>1.487</td><td>0.102</td><td>0.133</td><td>18.172</td><td>11.207</td><td>< 2e-16 ***</td><td><2e-16 ***</td></tr>
<tr><td>6 months</td><td>-0.622</td><td>-0.274</td><td>0.173</td><td>0.186</td><td>-3.594</td><td>-1.476</td><td>0.000326 ***</td><td>0.140</td></tr>
<tr><td>12 months</td><td>-0.468</td><td>-0.384</td><td>0.135</td><td>0.165</td><td>-3.461</td><td>-2.324</td><td>.000539 ***</td><td>0.020</td></tr>
<tr><td>18 months</td><td>-0.209</td><td>0.459</td><td>0.190</td><td>0.154</td><td>-1.102</td><td>2.981</td><td>0.270</td><td>0.00287 **</td></tr>
<tr><td>Post intervention</td><td>-0.116</td><td>-0.009</td><td>0.126</td><td>0.154</td><td>-0.921</td><td>-0.060</td><td>0.357</td><td>0.952</td></tr>
<tr><td>LLIN type</td><td>-0.235</td><td>-0.123</td><td>0.102</td><td>0.108</td><td>-2.297</td><td>-1.143</td><td>0.021611 *</td><td>0.253</td></tr>
</table>

Table 2: Generalised linear mixed model of *An. gambiae* s.l. and *An. funestus* s.l. vector density during the LLINEUP trial. Results of a nb-model <- glmmTB (species~ round+LLINtype+ (1|HH), family =n.bionm2). AG= *Anopheles gambiae* s.l.; AF= *Anopheles funestus* s.l.; HH= Household; n.bionm2= negative binomial type2, Std error= standard error; ***= p≤ 0.001; **=p ≤ 0.01 and *=p ≤ 0.05

Supplementary Figure 1

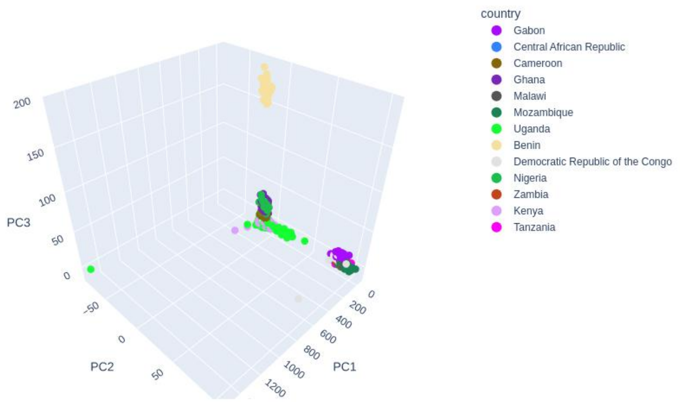


*Fig 1: An. funestus Principal component analysis.* *Using the genomic region 2R:60-80Mbps to avoid any polymorphic chromosomal inversions showing Ugandan LLINEUP An. funestus populations clustering with equatorial populations.*

Supplementary Figure 2

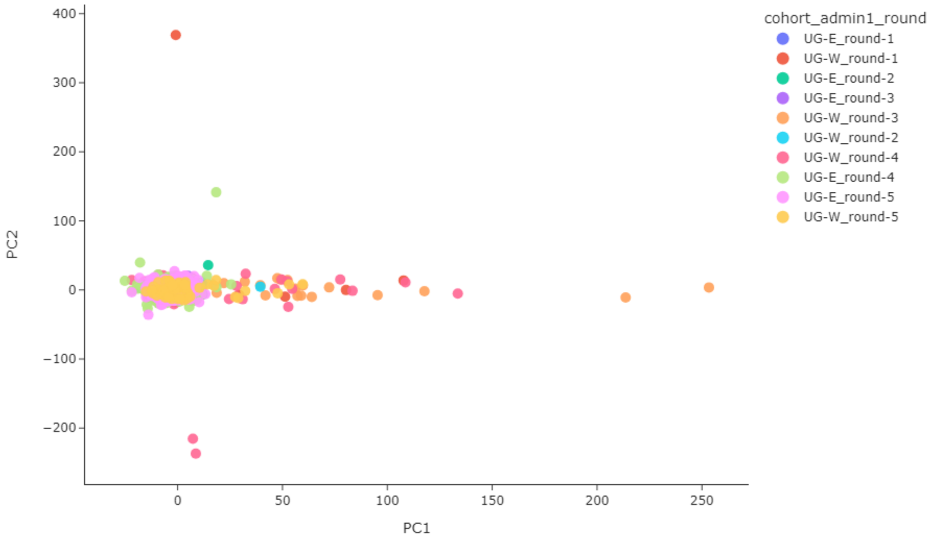


*Fig 2: An. funestus Principal component analysis.* *Using the genomic region 2R:60-80Mbps showing lack of structure within the An. funestus populations. Although geographically separated and collected over different temporal spaces, the East and Western Uganda An. funestus cluster together highlighting the lack of structure. UG-E= Uganda Eastern region, UG-W, Uganda Western region, round-1= Pre-intervention, round-2= 6 months, round-3= 12 months, round-4= 18 months, round-5= post- intervention.*

Supplementary Table 3

| Region | **Median $F_{ST}$ (*p value*)** |
|---|---|
| Between population | 0.00107 |
| Within West | 0.00306 |
| Within East | -0.00078 |
| Between population versus within West | 411754 (8.2718e-12) |
| Between population versus within East | 598502 (2.3845e-14) |

*Supplementary table 3: Pairwise $F_{ST}$ within East and Western Uganda versus between East and Western Uganda $F_{ST}$.*

Supplementary Figure 3

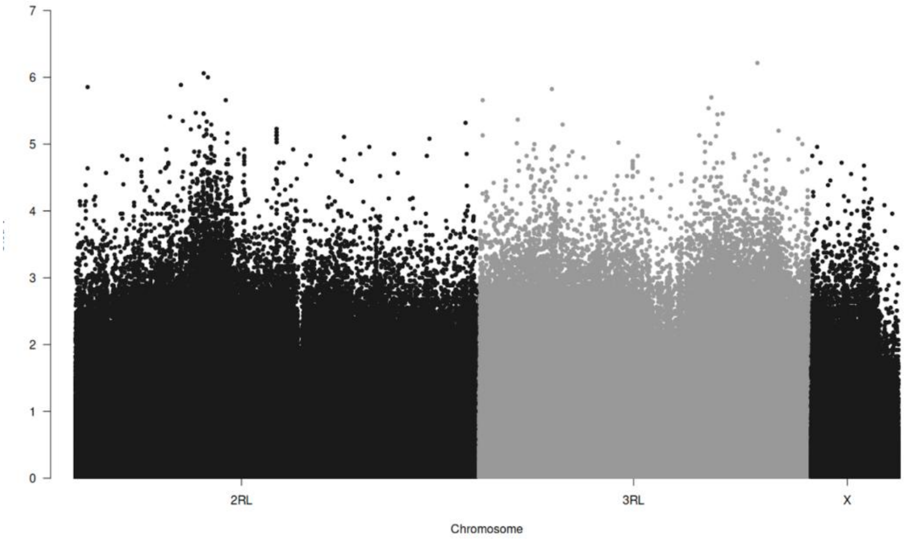


*Fig.3: Manhattan plot of SNPs between pre and post intervention An. funestus population. Used a generalised linear mixed model, and no SNP was significantly associated with LLIN use. Tested the association between allele frequencies and intervention (pre-intervention versus post-intervention) and net type as fixed effect, and health subdistricts as random effects.*

Supplementary Table 4:

| Chromosome | Gene | Peak center | Peak start | Peak stop |
|---|---|---|---|---|
| **2RL** | *RP1- locus* | 8667967 | 8658455 | 8683454 |
| | *Complexin transcript variant* | 51660985 | 51657957 | 51663230 |
| | | 57731386 | 57714091 | 57740033 |
| | Gste-2 | 67239558 | 67236023 | 67241326 |
| **3RL** | | 24915154 | 24855951 | 24944756 |
| | | 29245108 | 29237124 | 29249101 |
| | TMC-1 | 67929193 | 67925771 | 67931656 |
| **X** | *Cyp9k1* | 8452011 | 8448380 | 8459274 |
| | *Dgk* | 13639410 | 13636759 | 13641768 |

*Table 4: showing peak signal (p=0.05) detection in An. funestus. We identified regions within the genome that were under selection*

Supplementary Figure 4

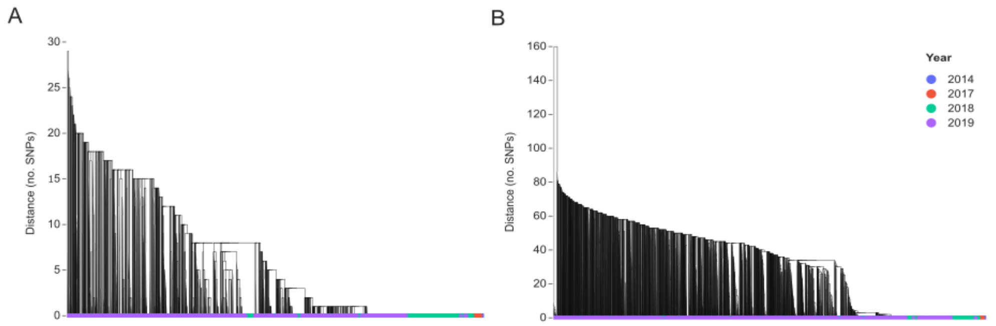


*Fig 4: Hierarchical clustering dendrogram of haplotypes over the TMC-1 (3RL: 67,925,771- 67,931,656- plot A&B) and Dgk (X: 13,590,000-13,690,044- plot C&D). Dendrogram leaves are labeled by year of sample collection (Plot A&B).*

Supplementary Figure 5

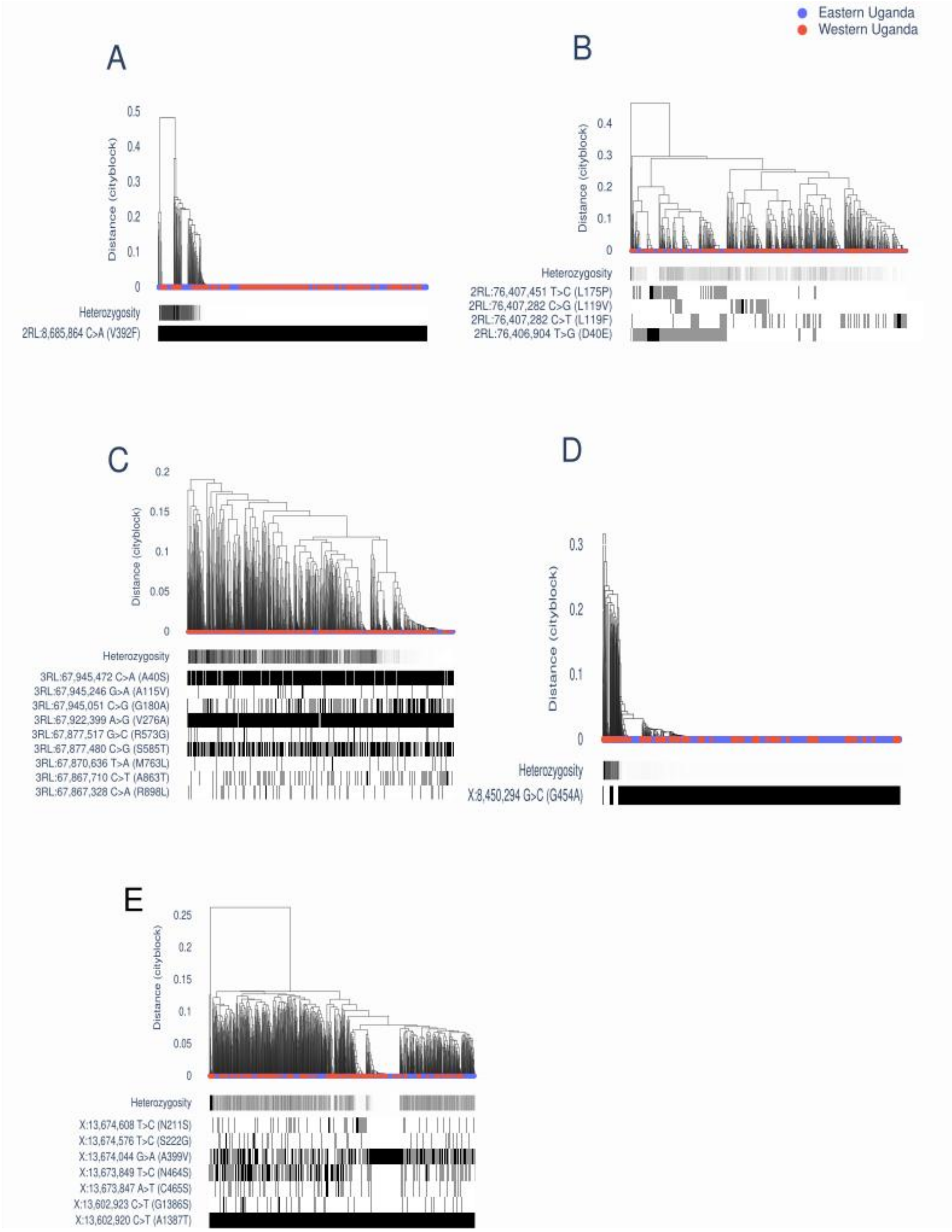


*Fig 5: Diplotype clustering: RP1 locus (2RL: 8,658,455-8,683,454-plot A), B) Gste2 (2RL: 76,406,705-76,407,655-plot B), TMC (3RL: 67,925,771-67,931,656-plot C), Cyp9k1 (X: 8,440,000-8,451,000-plot D) and Dgk (X: 13,590,000-13,690,044-plot E), and loci. Pairwise distance between diplotypes was calculated using genetic distance based on city-block distance and complete linkage plotted as hierarchical clusters. The colouring of the dendrogram leaves corresponds to the geographical*

*region from which each individual sample was collected; below the dendrogram is the assigned sample heterozygosity and gene SNP mutations are displayed as horizontal bars.*

Supplementary Table 5

| Region | Trial arm | Pre-intervention | 6 months | 12 months | 18 months | Post intervention |
|---|---|---|---|---|---|---|
| **Eastern** | Non-PBO | 3 | 45 | 17 | 230 | 195 |
| | PBO | 5 | 11 | 23 | 203 | 117 |
| **Western** | Non-PBO | 25 | 1 | 24 | 31 | 19 |
| | PBO | 6 | 1 | 2 | 48 | 39 |
| **Total** | | **39** | **58** | **66** | **512** | **370** |

*Table 5: Number of An. funestus collected from East and Western Uganda in both PBO and Non-PBO arms during the five rounds of the trial.*
